# Architecture of a DNA-guided Cas12a

**DOI:** 10.64898/2026.03.19.712971

**Authors:** Rodrigo Fregoso Ocampo, Carlos Orosco, Boyu Huang, Madeline S. West, Piyush K. Jain, David W. Taylor

**Author notes:** These authors contributed equally to this work (Rodrigo Fregoso Ocampo, Carlos Orosco).

## Abstract

CRISPR/Cas systems have largely been restricted to RNA-guided nucleases. Here, we present the cryo-EM structure of *Acidaminococcus* sp. Cas12a (AsCas12a) bound to a pseudo-DNA (ΨDNA) guide and RNA target, revealing how Cas12a accomplishes DNA-guided RNA recognition. The ΨDNA hairpin bridges the recognition and nuclease lobes, mimicking a PAM-proximal duplex and positioning the spacer to allow formation of a canonical RNA-DNA heteroduplex along the REC lobe. This provides a structural framework for its activity and provides a blueprint for future engineering.

---

CRISPR-Cas systems function as prokaryotic adaptive immune modules, utilizing sequence-guided nucleic acid target recognition. Beyond their biological role, these systems have been repurposed as versatile platforms for genome engineering in both fundamental research and therapeutic applications.^1,2^ CRISPR-Cas systems are canonically RNA-guided enzymes that direct programmable recognition of target sequences. Cas12a, a class 2 type V system, typically utilizes a CRISPR RNA (crRNA) guide that directs cleavage of double-stranded DNA targets containing a protospacer adjacent motif (PAM).^3,4^ Although several CRISPR nucleases have been adapted for RNA targeting,^5–8^ this activity generally relies on an RNA guide and possess non-specific collateral cleavage activity on bystander RNAs. Recent work from our group introduced a DNA-guided Cas12a repurposed to recognize RNA targets with high specificity using engineered ΨDNA guides.^9^ However, how Cas12a structurally accommodates a DNA guide while recognizing an RNA substrate remained unknown. Here, we determined the cryo-electron microscopy (cryo-EM) structure of AsCas12a bound to a ΨDNA guide and RNA target at 3.18 Å resolution, revealing how Cas12a accommodates an RNA target through structural mimicry of canonical CRISPR components (PDB: YYYY) (Figure 1a, Table 1, Supplementary Figure 1).

**Table 1.**
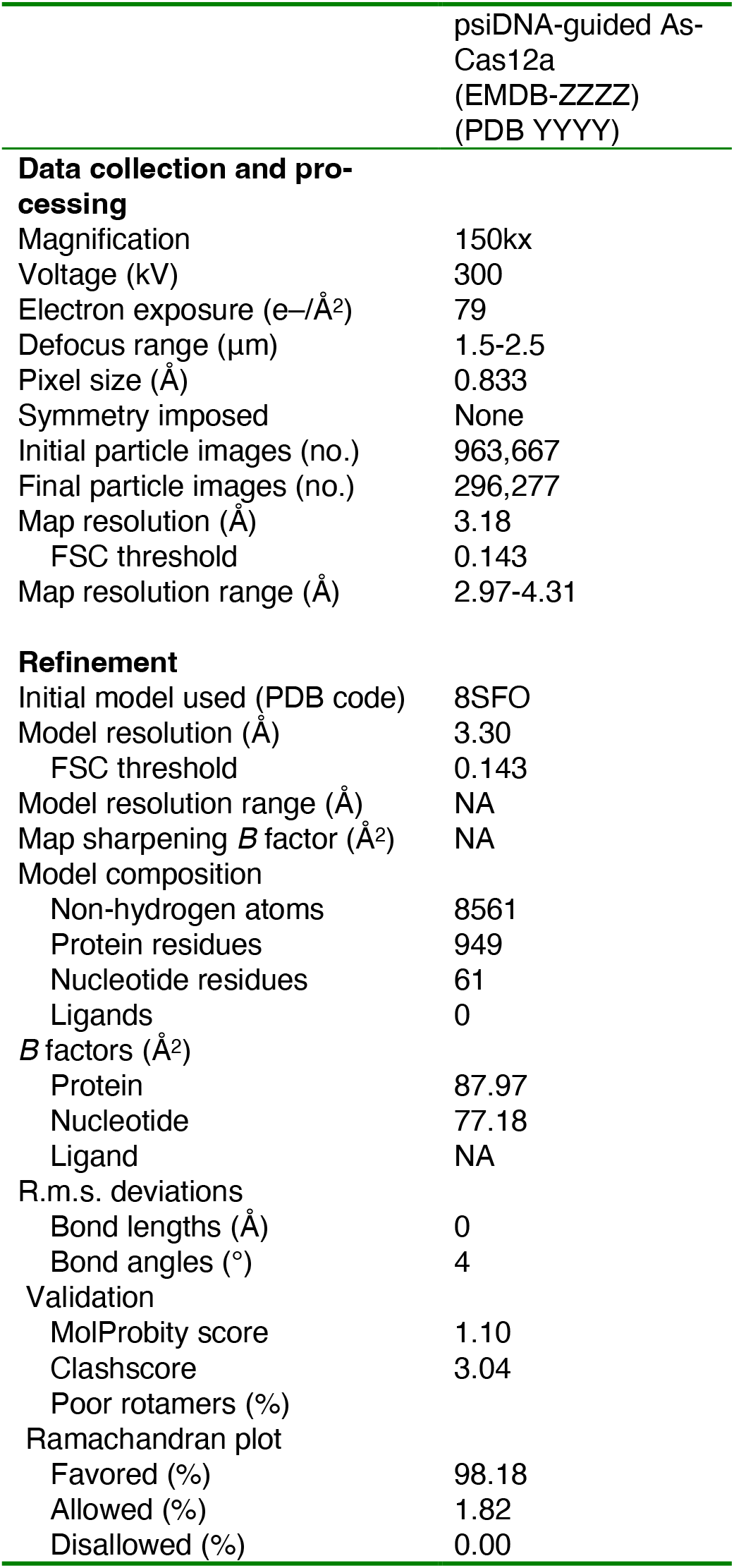
Cryo-EM data collection, refinement and validation statistics.

**Figure 1.**
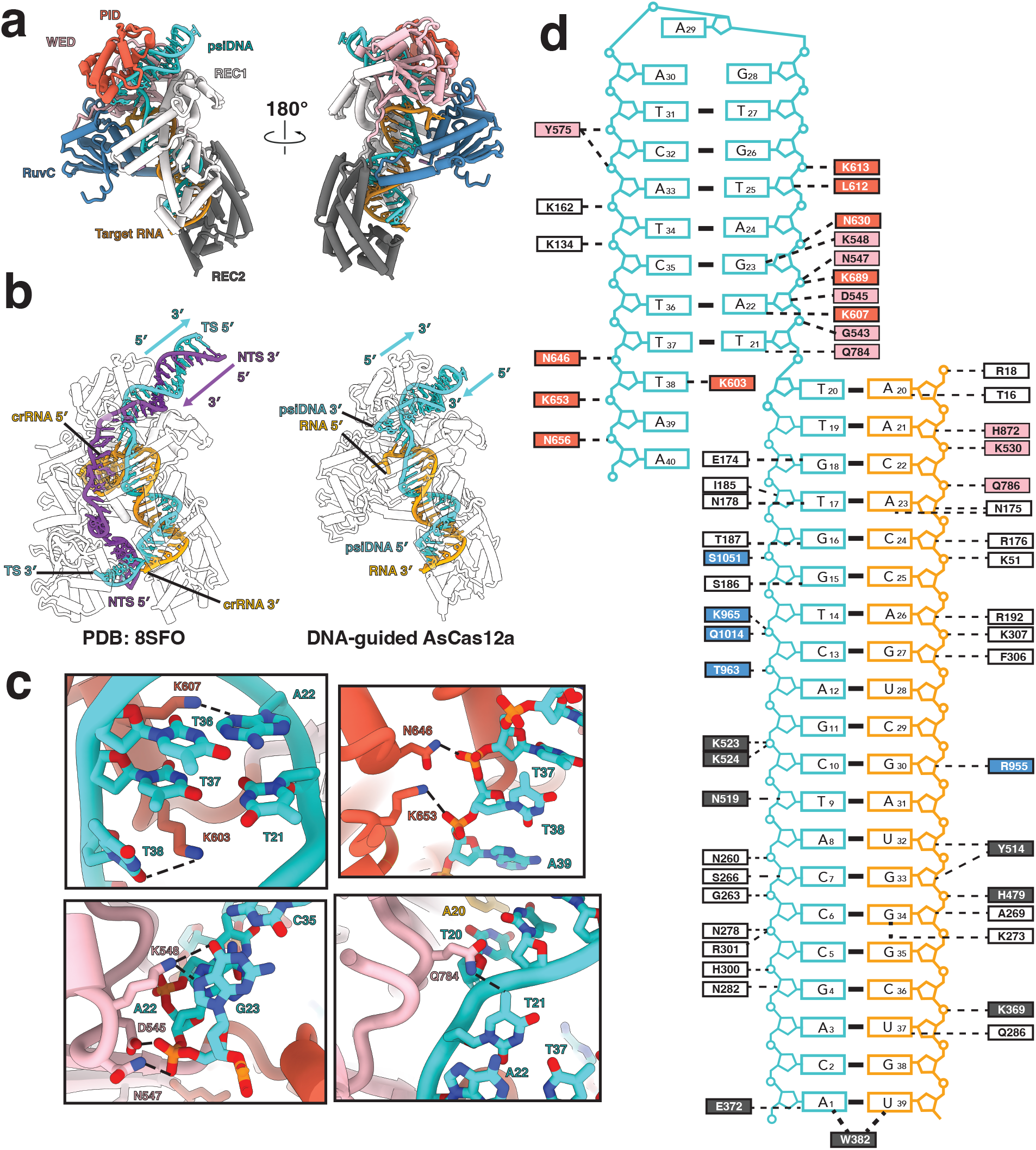
Structural basis for RNA targeting by ΨDNA-guided AsCas12a. **a**. Front and back views of the atomic model of ΨDNA-guided AsCas12a targeting a 40nt target RNA (PDB: YYYY). REC1 (snow), REC2 (dim grey), PID (tomato), WED (pink) and RuvC (steel blue) domains were resolved in the structure and modeled along with the ΨDNA (cyan) and target RNA (orange). b (Left) Atomic model of a 20bp AsCas12a effector targeting a canonical fully matched dsDNA duplex containing a canonical TTTN PAM (PDB 8SFO). The protein domains are transparent to allow visualization of the trajectory and directionality of both the TS (cyan), NTS (dark orchid), and crRNA (orange). (Right) AsCas12a atomic model bound to a ΨDNA targeting a 40 nt target RNA. The ΨDNA is bound in the same direction as the TS and NTS as recognized in the structure on the left. **c**. Visualization of PAM-analogous contacts formed between AsCas12a and the ΨDNA hairpin. K607 and Q784 form base-specific contacts with A22 and T37, respectively, while D545 and N547 in the WED domain form stabilizing contacts with the phosphate backbone of theΨDNA. In addition, K603 is intertwined with the end of the ΨDNA handle and contacts T38. N646 and K653 stabilize the phosphate backbone of the NTS-analogous end through non-specific contacts with the phosphate backbone **d**. Overview of the contacts formed between AsCas12a and the DNA:RNA heteroduplex. Many of the contacts to this heteroduplex are conserved including W382, which caps the PAM-distal end of the canonical R-loop after position 20. Notable contacts that are not formed are in the RuvC lid and bridge helix.

Our structure adopts the canonical bilobed architecture, resembling previously reported Cas12a structures in complex with crRNA and dsDNA targets.^10,11^ Surprisingly, the ΨDNA hairpin bridges the two lobes and binding the PID and WED domains in the same position that the PAM-proximal dsDNA helix would be recognized in RNA-guided systems (Figure 1A).^12^ The design of the ΨDNA, which contains a 3’-hairpin or handle and 5’-spacer, enables AsCas12a to bind the hairpin in the same directionality that it would bind a canonical dsDNA target duplex. This positions the ΨDNA spacer where the target strand would be in a canonical R-loop and with the same directionality (Figure 1b). This structure explains our recent findings that show that a 5’ handle is significantly less active than a 3’ handle, as this geometry of nucleic acids would be incompatible within this structural framework. In addition, the structure suggests that a dual handle ΨDNA with both a 5’ and 3’ hairpin would not significantly affect activity, as we reported. The 3’ handle would be properly recognized, the spacer would traverse through the enzyme, and the 5’ handle would be located outside of the core on the opposite end of the enzyme. Together, this indicates that the ΨDNA architecture uniquely recreates the spatial cues required for duplex recognition by an RNA-guided Cas12a.

Cas12a bends the ΨDNA hairpin to enable PAM-analogous contacts by amino acids previously characterized to recognize a 5’ TTTN PAM in a canonical dsDNA target duplex (Figure 1c).^13^ The ΨDNA contains residues that would correspond to a TCTT PAM. The presence of pyrimidines at positions –2 to –4, enable the hairpin to acquire a geometry like the one acquired by a canonical PAM. This enables AsCas12a to stabilize the hairpin utilizing base-specific contacts between T38 and K603 and G23 and K548, respectively, previously shown to be important in PAM recognition. The interaction between Cas12a and ΨDNA is further stabilized via a contact between Q784A and the final base of the duplex, T21, which forms a non-canonical base pair with T37 (Figure 1c). These interactions suggest that Cas12a recognizes the ΨDNA handle via a mechanism analogous to PAM binding but can tolerate deviations from canonical base pairing provided that the overall duplex geometry is preserved.

The structure also reveals a fully formed DNA:RNA heteroduplex scaffolded by the REC lobe of Cas12a (Figure 1d). This heteroduplex follows the same trajectory observed for an R-loop during canonical target recognition with the target RNA positioned where the crRNA is normally located. The non-complementary region of the 40-nt ssRNA target exits the structure at the interface of the WED and RuvC domains, where the crRNA hairpin would be expected to form in a canonical Cas12a structure. Since this region in the target RNA was designed to contain no secondary structure, it is not resolved in the cryo-EM reconstruction. The region of Cas12a that recognizes the crRNA hairpin was, likewise, not resolved in the structure. This flexibility is consistent with the absence of stabilizing interactions normally provided by the crRNA handle but consistent with previous reports that Cas12a can function with noncanonical crRNAs.^14^ These observations indicate that DNA-guided RNA targeting preserves the canonical hybrid nucleic acid-binding of Cas12a along the spacer region.

The DNA spacer region follows the standard trajectory that the DNA target strand would follow in the presence of a fully matched DNA duplex. This similarity suggests that the protein primarily recognizes the hybrid duplex through non-specific backbone interactions rather than strict nucleic acid identity. However, given that the ΨDNA is only 41 base pairs, the region that would be considered analogous to the non-target strand is incomplete, and therefore, only the first three bases (ΨDNA 38-40) are present in the structure. This region stabilizes the PAM-interacting domain by forming multiple contacts along its positively charged surface. These interactions partially compensate for the absence of the extended non-target strand normally present during DNA targeting. In a canonical AsCas12a structure, the rest of the non-target strand partitions through the RuvC and Nuc domains, enabling RuvC lid formation and R-loop maturation.^15,16^ However, our structure reveals that without a non-target strand, both the RuvC lid and the Nuc lobes are unresolved (Figure 2a). This observation indicates that stabilization of these domains depends on interactions with the displaced non-target strand. Because the RuvC lid is not present, multiple contacts with the TS, previously characterized in a 20bp R-loop complex, are not present in the structure, including stabilizing contacts by K965 and Q1013.^11^ Despite this, AsCas12a is still able to stably bind a fully formed double-stranded duplex with a fully docked bridge helix and REC2 domain, as evidenced by the presence of W382 capping the PAM-distal end of the R-loop.^17,18^

**Figure 2.**
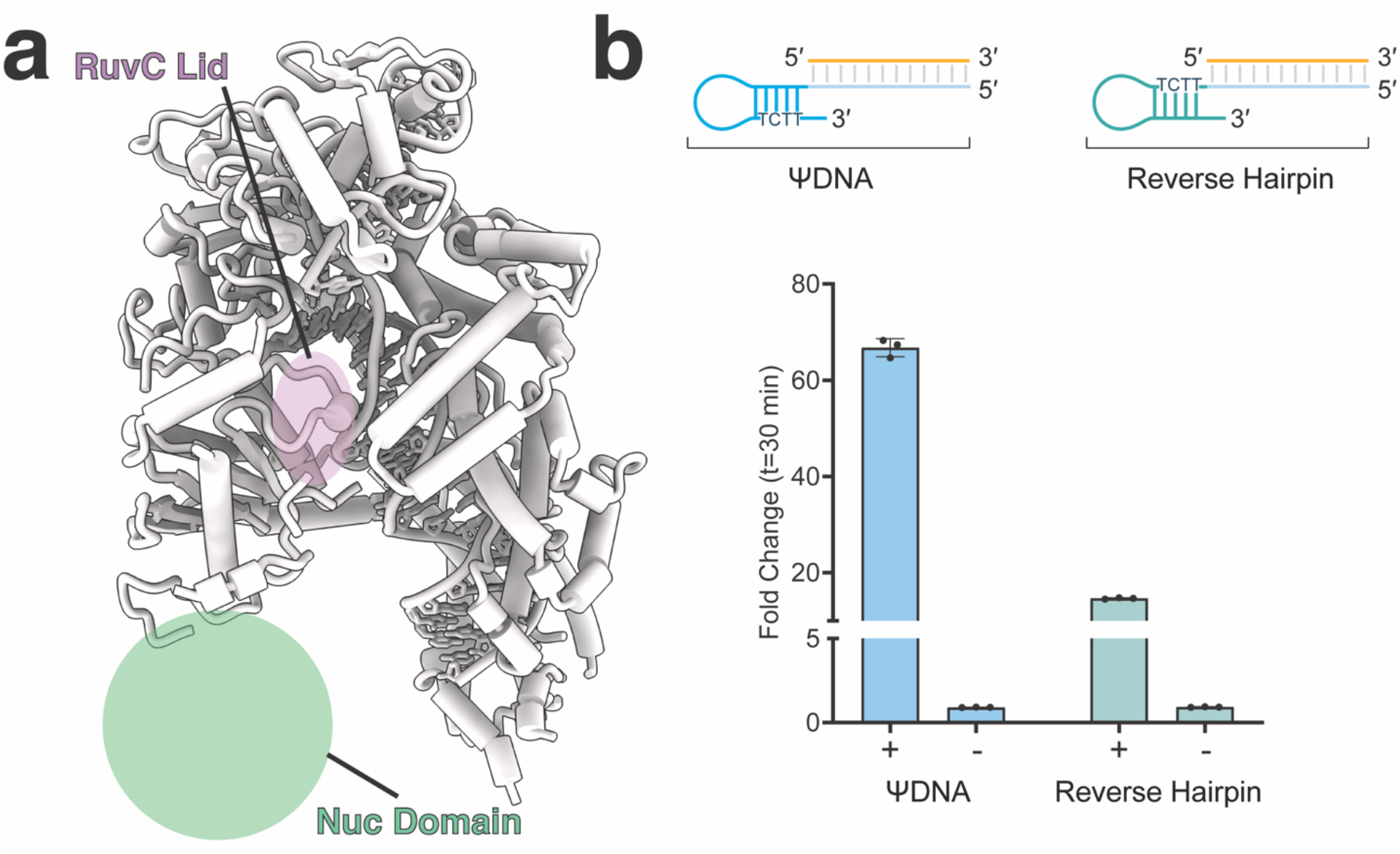
ΨDNA hairpin engineering can tune activity. **a**. The atomic model of ΨDNA-guided AsCas12a colored to depict two important missing regions in the structure. The Nuc domain (medium spring green) was not modeled in the structure due to high structural heterogeneity. The RuvC lid (plum) was also not resolved. **b**. Trans-cleavage assays using AsCas12a bound to a standard ΨDNA guide with a TCTT PAM and reverse hairpin ΨDNA guide containing a AGAT PAM. Fold change in RFU between final (t = 30 minutes) and initial (t = 0 minutes) reading for samples containing target RNA (+) or non-target RNA control (-). Error bars represent mean value +/- standards deviation (SD, *n* = 3).

It has been shown that PAM binding contributes to R-loop formation by enabling PAM-proximal bases of dsDNA to flip/twist to probe the sgRNA for complementarity.^19^ Since our ΨDNA design contains a ssDNA strand adjacent to the PAM and does not require this mechanism to occur to form a productive heteroduplex, we hypothesized that PAM recognition might not be strictly required in ΨDNA-guided AsCas12a, despite observing multiple conserved contacts. To test this, we flipped the bases of the hairpin of the ΨDNA to maintain hairpin stability, while simultaneously changing the region analogous to the PAM. This engineered PAM-analogous region was designed to contain a purine-rich AGAT sequence which diverges significantly from the TTTN PAM AsCas12a usually requires for activation. This ΨDNA design still showed significant trans-cleavage activity, despite a ∼3-fold reduction, indicating that ΨDNA can still accommodate a handle that is very different from the canonical PAM (Figure 2b). Together these results show that a pyrimidine-rich ΨDNA PAM-analogous region enhances trans-cleavage activity of Cas12a, but that Cas12a can still accommodate other sequences, if a stable hairpin is present. It is tempting to hypothesize that rationally designing changes to the ΨDNA sequence could lead to tunable activity.

Together, these findings reveal how Cas12a can accommodate DNA-guided RNA recognition through structural mimicry of canonical CRISPR nucleic acid components. Interestingly, these results are also consistent with a recent study that used AlphaFold to model DNA-guided Cas12a for RNA targeting.^20^ The ΨDNA hairpin recreates the geometry of the PAM-proximal duplex and is stabilized by interactions with the PAM-interacting domain, while the spacer region forms a canonical RNA-DNA heteroduplex within the REC lobe. The absence of a complete non-target strand alters the conformation of nuclease-domain elements associated with cleavage activation, highlighting the structural plasticity of Cas12a in accommodating alternative nucleic acid substrates. This structure provides a framework for understanding DNA-guided RNA-targeting Cas enzymes and offers insights that may guide the engineering of CRISPR systems for programmable RNA detection and manipulation beyond Cas12 systems.^14^ More broadly, this structural framework expands the potential targeting modalities of CRISPR nucleases.

## Methods

### Plasmid design and cloning

An AsCas12a protein was codon optimized for expression in E. coli and its DNA sequence was cloned into a pET28 expression vector under lacO repression with a 6x N-terminal His tag and a TEV protease using the NEBridge Golden Gate buffer set (NEB) and following manufacturer’s instructions.

### Oligonucleotide sequences

psiDNA:

ACAGCCCATCGACTGGTGTTTAGATGTGAATCATCTTTAAT

psiDNA Reverse hairpin

ACAGCCCATCGACTGGTGTTTTCTACTAAGTGTAGATTAAT

miR-21 RNA:

rGrArCrArUrUrUrArArCrArArCrArCrCrArGrUrCrGrArUrGrGrGrCrUrGrUrArGrCrArUrArA rArGrU

### Protein expression and purification

The cloned AsCas12a pET28 expression plasmid was transformed into BL21 DE3 chemically competent cells (NEB) and plated under kanamycin selection. A single colony was inoculated into 50 mL of Luria Broth (LB) under kanamycin antibiotic selection and was allowed to grow at 37 °C for 16 h. The inoculate was then back diluted 1:100-fold into two separate 1.5 L LB culture flasks and grown to OD600 0.8. Expression was induced by the addition of Isopropyl-β-D-thiogalactopyranoside (IPTG) at a final concentration of 0.1 mM and growth for 16 h at 18 °C to maximize expression levels. Cells were pelleted by centrifugation (4000 × *g*, 60 min) and resuspended using Lysis Buffer (20 mM tris-HCl pH = 7.5, 1 M NaCl, 10 mM MgCl2, 1 mM TCEP, 200 μM PMSF, 10% glycerol) with 1 Pierce Protease Inhibitor Tablet and 10 U DNAseI. Cells were lysed by sonication, and the lysate was clarified by centrifugation at 18,000 × *g* for 30 min at 4 °C. The supernatant was loaded into a 5 mL HisTrap HP Ni Column (Cytiva) and eluted using a gradient of Ni elution buffer (20 mM tris-HCl pH = 8.0, 500 mM NaCl, 250 mM Imidazole, 1 mM TCEP, 10% glycerol). Fractions containing the protein were combined and dialyzed with TEV protease in dialysis buffer (40 mM HEPES pH = 7.5, 150 mM NaCl, 1 mM TCEP, 10% glycerol) for 16 h at 4 °C. The sample was then purified by gel filtration chromatography (Superose 6 10/300; GE Healthcare) in dialysis buffer. The purity and quality of the protein were analyzed by SDS-polyacrylamide gels. The protein was quantified using the PIERCE Bradford assay reagent kit (Thermo Fisher) following the manufacturer’s instructions. The protein was flash-frozen in 6μL aliquots at a final concentration of 70.5 μM and stored at -80°C.

### In vitro trans-cleavage assay

The ΨDNA-AsCas12 trans-cleavage assays were prepared at 50 nM AsCas12a enzyme, 100 nM ΨDNA, 40 nM target RNA, and 500 nM FQ ssDNA reporter. All components were added in NEB 2.1 buffer and nuclease free water at a final volume of 40 μL. Then all reactions were plated on a dark flat-bottom 384-well plate (Grenier) and subsequently placed in a BioTek Synergy fluorescence microplate reader. Samples were incubated at a temperature of 37 °C for 30 minutes. Fluorescence was recorded at 483/20 nm and 530/20 nm excitation/emission wavelengths in 2.5-minute intervals.

### Cryo-EM sample preparation and data collection

For the 20-bp complex, a flash frozen sample of purified AsCas12a was rapidly thawed and incubated at a 10 μM final concentration with HPLC-purified ΨDNA containing a miR-21 targeting spacer at a 1:1 ratio at 37 °C for 30 minutes. After a 30-minute incubation a 40-nt target RNA containing a region complementary to the miR-21 targeting spacer was added at a final concentration of 10μM and further incubated for 30 minutes at 37°C. The sample was then allowed to cool on ice before being applied to Quantifoil 1.2/1.3 Ultra Thin 400-Mesh Carbon Grids that had been glow discharged with 20 mA for 30 s prior to blotting. All grids were blotted using a Vitrobot Mark IV (Thermo Fisher) for 9-12 s, blot force set to 1 at 4 °C and 100% humidity, and plunge-frozen in liquid ethane. Data were collected on an FEI Titan Krios 300 V Transmission Electron Microscope (Thermo Scientific) with a Gatan BioContinuum Imaging Filter and a K3 direct electron detector. Data were collected on Serial EM v4.2 with a 0.832 Å pixel size and a defocus range of – 1.5 to –2.5 μm with a total exposure time of 3.3 s in a total accumulated dose of 79e/A^2^ at -30° tilt. Motion correction, contrast transfer function (CTF) estimation, and particle picking were carried out in cryoSPARC live v4.7.1 4338 movies were accepted for downstream processing.

### Cryo-EM image processing, model building and refinement

From the accepted movies, a total of 1,767,373 particles were selected using Blob Picker. A total of 1,107,909 particles were picked after a single round of 2D classification. A random subset of 100,000 picked particles was selected for ab-initio reconstruction (3 classes). The best class was used for two rounds of heterogeneous refinement and downstream non-uniform refinement, yielding a 3.10 Å reconstruction comprising 963,667 particles. Given that the Nuc domain was mostly unresolved, particle homogeneity was enhanced by performing a single round of 3D classification. All classes contained either a partially resolved or fully unresolved Nuc domain. Classes containing the fully unresolved Nuc domain were selected for a final non-uniform refinement, yielding a final 3.18Å reconstruction comprised of 296,227 particles

### Model Building and Refinement

Since the structure resembled a fully formed 20bp R-loop conformation, the initial backbone for the model was built using a previously solved structure (PDB: 8SFO) and aligning it to the map. The alignment yielded a well-aligned DNA:RNA heteroduplex as well as the alpha helices from the protein. The output was used as a fiducial for de-novo model building using a combination of coot v1.0 and ChimeraX v1.90 The DNA non-target strand was deleted from the map as well as the Nuc domain. The DNA hairpin was modeled manually using Isolde v1.4. All side chains rotamer optimization and Ramachandran angle refinement were performed using a combination of coot v1.0 and Isolde v1.4. Multiple iterations of real space refinement implemented within Phenix v1.19 were carried out to improve the model geometry and map fit

## Data availability

All the data supporting the findings of this study are available within the Article and Supplementary Files. Additional data can be obtained from the corresponding authors, D.W.T. or P.K.J., upon reasonable request. The atomic coordinates have been deposited in the Protein Data Bank under accession code PDB YYYY. The cryo-EM density map has been deposited in the Electron Microscopy Data Bank under accession code EMD-ZZZZ.

## Acknowledgements

We extend our gratitude to the University of Texas Austin and the University of Florida (UF) Health Cancer Center for their support. We also thank the members of the Taylor and Jain labs for valuable discussions and for their help with experiments. This work was funded by the Exxon Mobil Gator Alumni Faculty and Shah Foundation Endowment Funds (P.K.J.), NIH-NIAID R61AI181016 (P.K.J.), NIH-NIGMS R35GM147788 (P.K.J.), and NIH-NIGMS R35GM158551 (D.W.T.). Computational resources for this work were supported by Welch Foundation grant F-1938 (D.W.T.). The funding sources had no role in study design, data collection, analysis, interpretation, or manuscript preparation.

## Author information

These authors contributed equally to this work: Rodrigo Fregoso Ocampo and Carlos Orosco.

### Contributions

R.F.O. and M.W. performed cryo-EM structure determination, model building, and structural analysis. R.F.O. and C.O. purified and reconstituted the enzyme complexes, while C.O. and B.H. conducted biochemical experiments. Data analysis and interpretation and manuscript preparation were performed by R.F.O., C.O., P.K.J., and D.W.T. P.K.J. and D.W.T. supervised the study and secured funding.

