## Supplementary Information for "Architecture of a DNA-guided Cas12a"

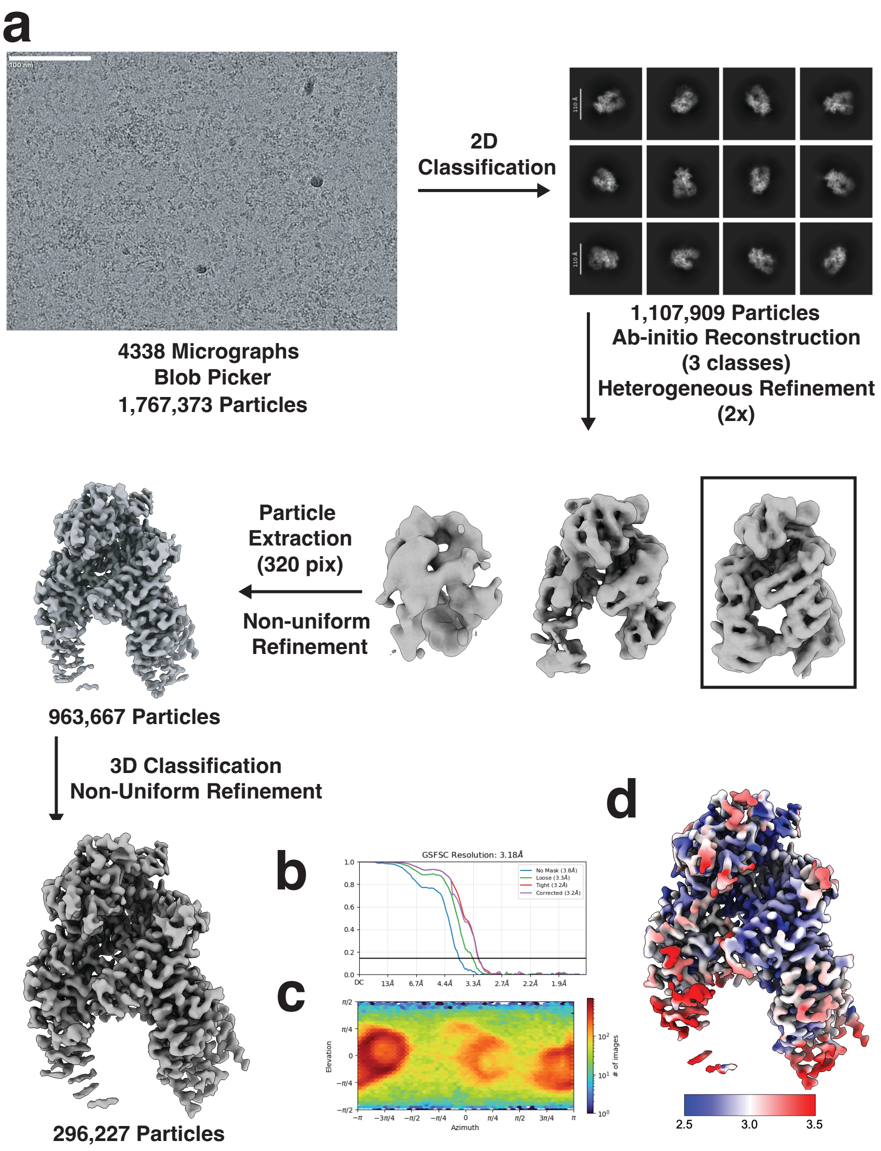


**Supplementary Fig. 1. Cryo-EM dataset processing pipeline, data analysis and quality control a.** Processing pipeline for ΨDNA-guided RNA targeting AsCas12a. **b.** Fourier Shell Correlation (FSC) diagram for the final cryo-EM structure obtained. **c.** Euler particle distribution diagram shows a well-represented orientation distribution of particles for the final cryo-EM structure. **d.** Sharpened map colored by local resolution with gold-standard FSC at a threshold of 0.143.
